# The SARS-CoV-2 Omicron sub-variant BA.2.86 is attenuated in hamsters

**DOI:** 10.1101/2023.11.10.566576

**Authors:** Vanessa Herder, Diogo Correa Mendonca, Nicole Upfold, Wilhelm Furnon, Karen Kerr, Georgios Ilia, Jay Allan, Alex Sigal, Arvind H. Patel, Massimo Palmarini

## Abstract

SARS-CoV-2 variants have emerged throughout the COVID-19 pandemic. There is a need to risk-assess newly emerged variants in near “real-time” to estimate their potential threat to public health. The recently emerged Omicron sub-variant BA.2.86 raised concerns as it carries a high number of mutations compared to its predecessors. Here, we assessed the virulence of BA.2.86 in hamsters. We compared the pathogenesis of BA.2.86 and BA.2.75, as the latter is one of the most virulent Omicron sub-variants in this animal model. Using digital pathology pipelines, we quantified the extent of pulmonary lesions measuring T cell and macrophage infiltrates, in addition to alveolar epithelial hyperplasia. We also assessed body weight loss, clinical symptoms, virus load in oropharyngeal swabs, and virus replication in the respiratory tract. Our data show that BA.2.86 displays an attenuated phenotype in hamsters, suggesting that it poses no greater risk to public health than its parental Omicron sub-variants.

**Article summary line:** The newly emerged Omicron sub-variant BA.2.86 is attenuated in hamsters.

## Main text

The coronavirus disease-19 (COVID-19) pandemic is characterised by the continuous emergence of Severe Acute Respiratory Syndrome Coronavirus-2 (SARS-CoV-2) variants (1). The Omicron subvariant BA.2.86 was initially detected from samples collected in July 2023, and subsequently designated as a variant under monitoring (VUM) by the World Health Organisation after its detection in multiple countries (2). BA.2.86 is most closely related to BA.2 circulating in Southern Africa in early 2022 (3). The concerns related to BA.2.86 emergence are due to the high number of mutations compared to its predecessor (50 along the entire genome, including 33 in spike) (4). BA.2.86 does not appear to escape existing population immunity (conferred by either XBB.1.5, or BA.4/BA.5 breakthrough infections) significantly more than the currently dominating variants XBB.1.5 and EG.5.1 (4). A recent report from the UK, also suggests that individuals that have not been vaccinated since 2022 maintain sufficient cross-neutralising activity against BA.2.86 (5).

In addition to their immunoevasive properties, emerging new variants need also to be risk-assessed for their intrinsic virulence. The BA.1 variant displays an attenuated phenotype, compared to the previous circulating variants (6). However, there is no evolutionary pressure on the virus to decrease virulence, unless this affects virus transmission.

Hamsters have proved to be an excellent model to assess SARS-CoV-2 virulence (7) as they are susceptible to SARS-CoV-2 and develop lesions in the respiratory tract resembling those induced in humans. Importantly, so far there has been a good correlation between the virulence of SARS-CoV-2 variants in hamsters and humans. For example, the higher level of virulence of Delta in the human population, compared to the original B.1, has been recapitulated in hamsters (8).

We have recently developed a digital pathology pipeline to quantify SARS-CoV-2-induced lesions in the respiratory tract, as a proxy for viral virulence (9). Using these methods, we confirmed how Delta is more virulent than BA.1. We also showed that Omicron sub-variants BA.2.75 and the recently emerged EG.5.1 have regained some virulence, although not reaching the levels observed for the pre-Omicron variants (9).

Here, we experimentally infected hamsters with either BA.2.86 (n=8), BA.2.75 (n=6), or mock-infected controls (n= 4). We compared BA.2.86 virulence to BA.2.75, as in our hands the latter is among the most virulent of the Omicron sub-variants. We monitored animals’ weight and clinical signs daily until 6 days post-infection (dpi), before culling them for post-mortem analyses. Tissues were fixed in formalin and embedded in paraffin wax as described previously (9). We also took oropharyngeal swabs for RNA extraction and quantification of viral load at 1, 2 and 5 (dpi). Animals infected with BA.2.75 displayed a moderate weight loss, while animals infected with BA.2.86 gained some weight during the duration of the experiment, although not as pronounced as in the animals of the mock-infected group (Fig. 1A). All (6/6) BA.2.75-infected, as opposed to only 3/8 BA.2.86-infected, hamsters displayed mild clinical signs (Fig. 1B). The remaining 5 BA.2.86-infected animals, like the mock-infected group, showed no clinical signs. Virus load was ∼ 3-7 fold higher in BA.2.75-infected animals, compared to those infected with BA.2.86 at 1 and 2 dpi, while virus RNA was detectable at 5 dpi only in 5/6 BA.2.75-infected animals (Fig. 1C). We quantified virus-induced pulmonary lesions using software-assisted whole lung section (collected at 6 dpi) imaging of immunohistochemistry, and downstream automatic quantification of positive signal, as previously described (9). Briefly, we quantified the levels of alveolar epithelial hyperproliferation using TTF1 antibodies, but excluding by machine learning approaches individual TTF1^+^ type 2 pneumocytes and epithelia bronchial cells (Fig. 1D). We quantified T cell and macrophage infiltrates using CD3 (Fig. 1E) and IBA1 (Fig. 1F) antibodies, respectively. We detected higher levels of alveolar epithelial proliferation, T-cell and macrophages infiltrates in the lungs collected from BA.2.75 infected hamsters, while lungs from BA.2.86-infected animals were not significantly different from mock-infected controls (Fig. 1G-I).

**Figure 1.**
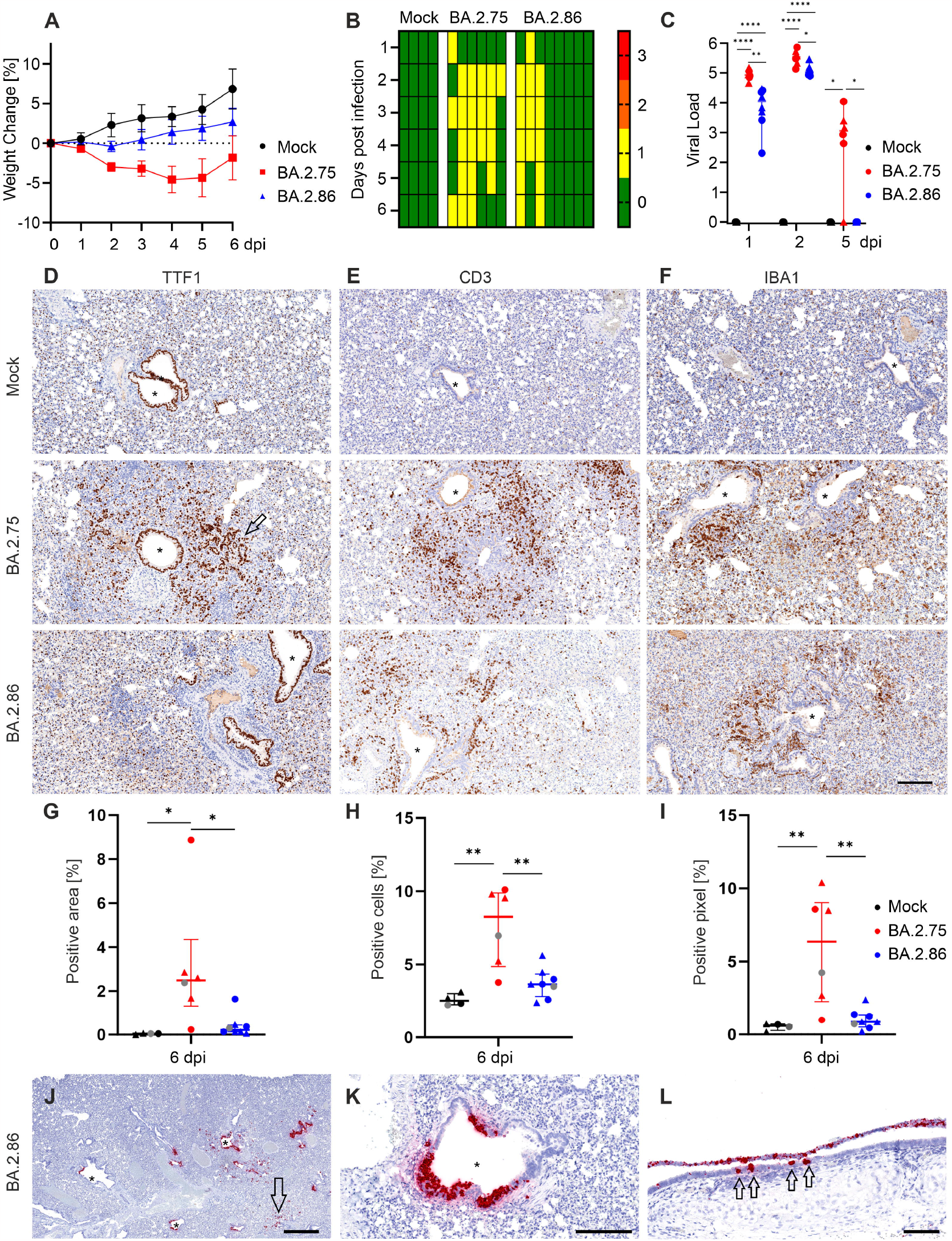
Virulence of the Omicron BA.2.75 and BA.2.86 sub-variants in experimentally infected hamsters. Hamsters were infected intranasally with BA.2.75 or BA.2.86 (50 microlitres of 3.75×10^6^ genome copies equivalent per animal) or mock-infected. A) Weight changes of hamsters are provided as mean with standard deviation. B) The clinical scores of hamsters are shown for the time period of 6 dpi. The disease scores of each animal was recorded daily. 0 – No weight loss or clinical signs observed. Values are as follows: 1 = small weight loss (under 15%) and mild clinical signs; 2 = weight Loss (15-20%) and noticeable clinical signs. 3 = weight Loss (over 20%) and severe clinical signs. C) Viral load obtained by total RNA extraction from throat swabs eluted in DMEM (Thermo Fisher) using the Direct-zol™-96 MagBead RNA Kit (Zymo Research; R2102), and SARS-CoV-2 genomic RNA was quantified by RT-qPCR as previously described (9). Data are shown as log_10_ genome copies/β-actin. Two-way ANOVA, Tukey’s post-hoc test; scatter dot plot, median with range. Significance is indicated with ^*^<0.05, ^**^<0.01, ^***^<0.001, ^****^<0.0001. D) Immunohistochemistry of lung sections stained with TTF1 shows proliferation of type-2-pneumocytes (T2P), and a positive signal in the nuclei of normal bronchi (*). Mock infected animals (top row), in BA.2.75-infected animals (middle row), the arrow highlights the proliferation of T2P in the parenchyma. The bottom row shows animals infected with BA.2.86. E) T cell infiltrates detected by immunohistochemistry using CD3 antibodies. In mock-infected lungs, T cells are scattered around in the lung parenchyma (top row), while in lungs of hamsters infected with BA.2.75 (middle row), and in BA.2.86-infected hamsters (bottom row) a diverse degree of infiltration of T cells is visible in the lung parenchyma around bronchi (*). F) Infiltrates of alveolar macrophages are present in mock infected animals (top row), severe in the BA.2.75 infected animals (middle row) and mild in BA.2.86 infected hamsters (bottom row). The asterisk (*) indicates bronchial epithelium. Scale bar is representative for all pictures, 200 micrometres. (G-I) Software-assisted quantification of lung pathology. G) BA.2.75-infected hamsters show significantly more proliferation of T2P compared to mock and BA.2.86-infected hamsters. H) Significantly more T cells (CD3) have been detected in BA.2.75-infected hamsters compared to mock and BA.2.86. I) Quantification of IBA1^+^ pixel in the lungs of BA.2.75 infected hamsters is significantly higher compared to mock and BA.2.86-infected hamsters. Note that grey data points in G-I represent the values of the animals displayed in the photomicrographs above. One-way ANOVA, Tukey post-hoc test; scatter dot plot, median with range. Significance is indicated with ^*^<0.05, ^**^<0.01, ^***^<0.001, ^****^<0.0001. Male animals: triangles, female animals: circles. Male animals are shown in triangles, female animals are depicted in circles. J) Detection of viral RNA in the lungs of hamsters infected with BA.286at 2 dpi. Bronchi (*) show a positive signal for viral RNA (signal in red) while in the parenchyma only minimal signal was detected (arrow). Bar, 1 mm. K) In the higher magnification, the bronchial epithelial cells show a strongly positive signal (red) in the cytoplasm, often positive cells are clumping together. Bar, 200 micrometres. L) The luminal trachea shows a slough-off of virus positive cells (top layer) as well as positively stained (red) epithelial cells (arrows). Bar, 100 μm. Images of the whole organ tissue sections shown above are available online at the CVR virtual microscope (https://covid-atlas.cvr.gla.ac.uk/). Raw data used to generate the data illustrated in this figure are available in Enlighten at the following doi: 10.5525/gla.researchdata.1528

We also infected an additional groups of hamsters with BA.2.86 (n=4), and culled them at 2 dpi in order to assess viral replication in the lungs at the early times post-infection, as we and others have shown that virus is largely cleared by 6 dpi (9). As expected, in BA.2.86 infected animals we detected virus in the trachea, as well as large bronchi and only very occasionally in the lung parenchyma (Figure 1J-L).

The results presented above in experimentally infected hamsters, are in line with a recent observation in England, where 45 infections caused by BA.2.86 were diagnosed in residents and staff of a care home (10). Although BA.2.86 showed a very high attack rate (87%), less than half of the infected patients were symptomatic and only one was hospitalised. Overall, our study shows that the newly emerging variant BA.2.86 is relatively non-pathogenic in hamsters. Our data suggest that at present, this variant should not pose a greater risk than the original Omicron variant.

## Acknowledgements

The authors thank the Histology Research Service, University of Glasgow for their excellent support and the high quality of their services. We thank Derek Gatherer for maintaining the online resource “CVR Virtual Microscope”.

## Contribution

Investigation : VH, DCM, NU, WF, KK, GI, JA; Formal Analysis: VH, DCM, NU; Conceptualisation: VH, AHP, MP; Data curation: VH, DCM, NU; Funding Acquisition: AHP, MP; Resources: AS; Project administration: VH, AHP, MP; Supervision: VH, AS, AHP, MP; Visualisation: VH, DCM, NU; Writing original draft: VH, WF, MP; Writing – editing and review: all.

